# Restoring local climate refugia to enhance the capacity for dispersal-limited species to track climate change

**DOI:** 10.1101/2022.07.25.501473

**Authors:** Gregory A. Backus, Christopher F Clements, Marissa L. Baskett

**Affiliations:** University of California, Davis; University of Bristol

**Keywords:** climate change, climate refugia, dispersal, environmental stochasticity, heterogeneity, metapopulations, population modeling, restoration

## Abstract

Climate refugia are areas where species can persist through climate change with little to no movement. Among the factors associated with climate refugia are high spatial heterogeneity, such that there is only a short distance between current and future optimal climates, as well as biotic or abiotic environmental factors which buffer against variability in time. However, climate refugia may be declining due to anthropogenic homogenization of environments and degradation of environmental buffers. To quantify the potential for restoration of refugia-like environmental conditions to increase population persistence under climate change, we simulated a population’s capacity to track increasing temperatures over time given different levels of spatial and temporal variability in temperature. To determine how species traits affected the efficacy of restoring heterogeneity, we explored an array of values for species’ dispersal ability, thermal tolerance, and fecundity. We found that species were more likely to persist in environments with higher local heterogeneity and lower environmental stochasticity. When simulating a management action that increased the local heterogeneity of a previously homogenized environment, species were more likely to persist through climate change, and population sizes were generally higher, but there was little effect with mild temperature change. The benefits of heterogeneity restoration were greatest for species with limited dispersal ability. In contrast, species with longer dispersal but lower fecundity were more likely to benefit from a reduction in environmental stochasticity than an increase in spatial heterogeneity. Our results suggest that restoring environments to refugia-like conditions could promote species’ persistence under climate change in addition to conservation strategies such as assisted migration, corridors, and increased protection.

## Introduction

Many species are tracking climate change by shifting their ranges to higher latitudes and elevations (Parmesan et al. 1999, Chen et al. 2011). Species that are adapted to a narrow range of climate conditions that cannot shift to analogous climates quickly enough could be at risk of extinction (Pearson 2006, Urban 2015). Traits that could limit a species’ ability to track climate change include short dispersal, narrow climate tolerance, or low fecundity (Pearson 2006, Urban 2015). The ability to track climate change also depends on characteristics of the environment, as variation in spatial heterogeneity and topography mean that local climate will change at different speeds in different areas across multiple scales (Ashcroft et al. 2009).

Areas that are expected to face relatively limited climate change represent “climate refugia” where species might persist with little to no movement (Dobrowski 2011, Morelli et al. 2016). Paleoecological research suggests that many dispersal-limited tree species survived past rapid climate change events in climate refugia because physical geographic characteristics, such as topography, buffered against rapid shifts in climate (Gavin et al. 2014). Though climate refugia often arise from physical characteristics such as topography (Dobrowski 2011), snowmelt (Millar et al. 2015), hydrology (McLaughlin et al. 2017), or upwelling in coral reefs (Chollett & Mumby 2013), climate refugia could also be the result of biological characteristics such as community composition and canopy cover (Lloret et al. 2012; De Frenne et al 2013).

Preserving climate refugia could support the persistence of climate-vulnerable populations (Morelli et al. 2016). This management approach holds potential for species that already live in areas that already contain climate refugia, which could include many climate-vulnerable species (Harrison & Noss 2017). However, because many common methods for identifying climate refugia rely on paleoecological records that do not reflect recent landscape changes, they may have limited utility in making predictions for modern climate change (Keppel et al. 2012). In particular, human development in biodiversity hotspots and climate refugia could reduce the availability of climate refugia for species that live within them (Cincotta et al. 2000). Climate-vulnerable species that do not currently live near or within climate refugia might then require alternative conservation strategies (Roberts & Hamann 2016), such as restoring the environmental conditions that might have previously supported refugia.

One potential source of climate refugia that could be impacted by anthropogenic landscape change is heterogeneity in climatic conditions over space (Hampe & Jump 2011; Ashcroft et al. 2012). High climate heterogeneity can reduce the distance a species needs to move to reach analogous habitats, allowing species to track climatic change by dispersing locally rather than dispersing across large latitudinal distances (Fig. 1e; Brito-Morales et al. 2018). However, anthropogenic homogenization of landscapes has decreased biodiversity across terrestrial (McKinney & Lockwood 1999; Groffman et al. 2014), freshwater (Scott & Helfman 2001), and marine (Thrush et al. 2006) ecosystems, leading to conservation proposals to restore local-scale heterogeneity and restore the natural processes that historically supported biodiversity in these locations (Palmer et al. 2010; Morelli et al. 2016). Restoring heterogeneity might therefore buffer against the effects of climate change, but restoring climate heterogeneity might negatively impact some species, such as those with niche specialization in the microclimate most prevalent in the homogenized environment (Allouche et al. 2012).

**Figure 1:**
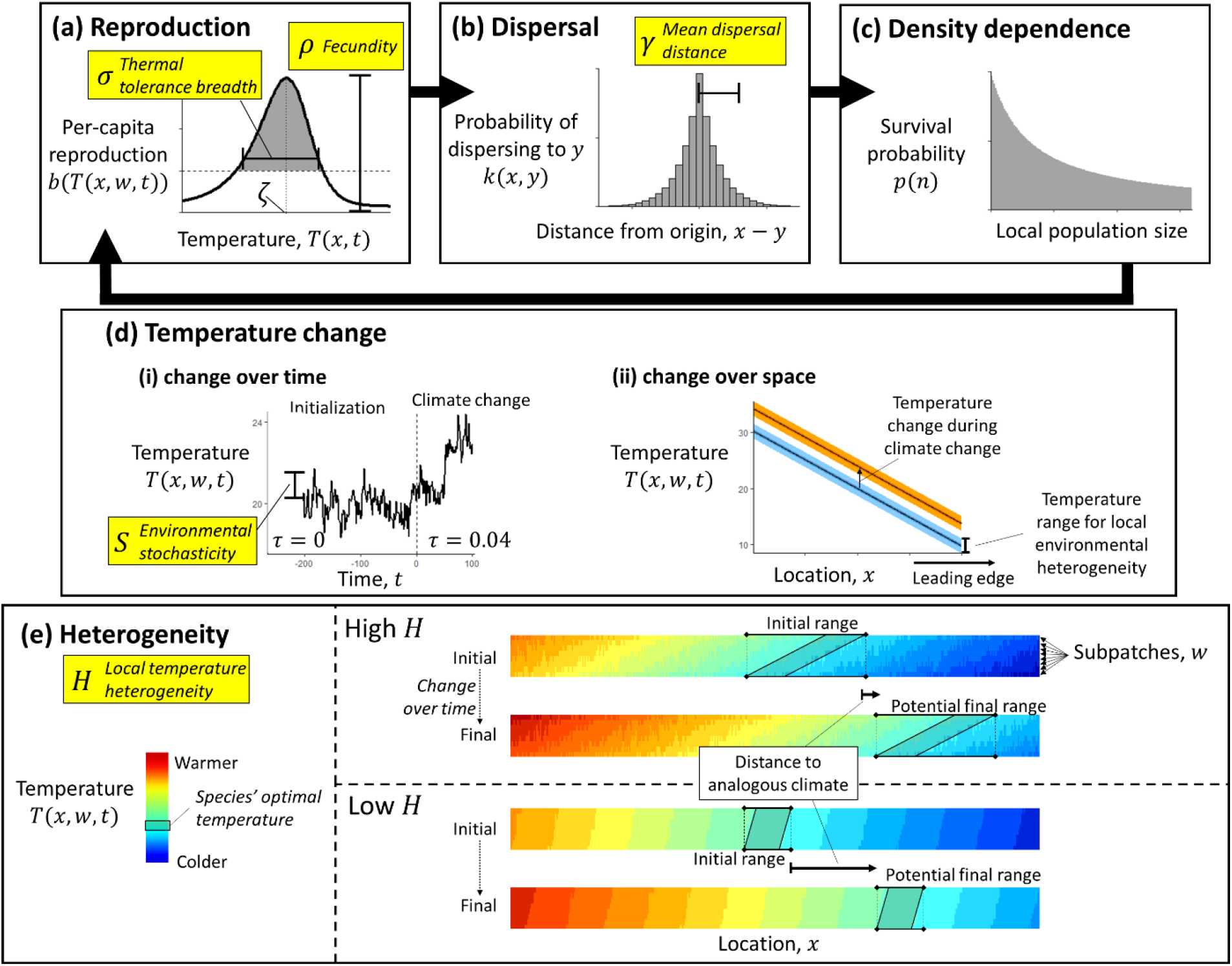
During each time step of the model, the population cycles through (a) reproduction, (b) dispersal, and (c) density dependence, then (d) the mean temperature changes, determining the environmental conditions for the next time step. We explore a range of values for parameters highlighted in yellow. (a) Per capita reproductive output *bi*(*T*(*x, w, t*)) is skew-normal, dependent on temperature *T*(*x, w, t*). This function depends on species’ thermal optimum *ζ*, thermal tolerance breadth *σ*, and fecundity *ρ*. (b) Individuals disperse across patches with a mean dispersal distance *γ*. After arriving in the destination patch, they redistribute among local subpatches. (c) Individuals compete over limited space, where each subpatch has a carrying capacity *K*. In each subpatch, individual survival probability *p*(*n*) decreases as local population size increases. (d.i) Temperature changes over time depending on environmental stochasticity parameter *S*. After an initialization phase with no temperature change (τ=0.04), the model shifts to the climate change phase (*τ* = 0.04). (d.ii) Mean temperature decreases linearly over space from the equatorward to the poleward side, and each location *x* has a range of temperatures from local heterogeneity between subpatches. During climate change, the average temperature increases by 4°C (or by 2°C) over 100 years. (e) Each patch in space contains *W* subpatches. The standard deviation of the temperature among subpatches within a patch depends on the local temperature heterogeneity *H*. Between the beginning and end of climate change, analogous temperatures are closer with higher rather than lower local temperature heterogeneity.

Another source of climate refugia impacted by anthropogenic activities are processes that buffer local interannual climate variability (Hampe & Jump 2011). Protected species’ distributions are often more accurate at predicting suitable conditions for species when they account for interannual variation (Zimmermann et al. 2009; Briscoe et al. 2016). In future projections, higher environmental stochasticity increases the likelihood of population extinction (Lande 1993), and higher interannual variation and frequency of extreme weather events during climate change could increase this extinction risk (Jentsch et al. 2007; Wernberg et al. 2013). Anthropogenic disturbance in locations with properties that buffer interannual climate variation could then reduce the availability of critical climate refugia, as might anthropogenic impacts that affect abiotic or biotic buffers (e.g., impacts that reduce canopy cover; De Freene et al. 2013).

Here, we quantify the potential for restoring human-altered environments to past levels of spatial heterogeneity or environmental stochasticity to create climate-refugia-like conditions that promote species persistence under climate change. Specifically, we use a metapopulation model to explore how local climate heterogeneity and environmental stochasticity affect a species’ ability to persist through climate change. The model cycles through reproduction, dispersal, and density dependence, where variation in temperature from spatial heterogeneity, environmental stochasticity, and climate change affect reproduction. By exploring a range of values for biotic parameters (dispersal distance, thermal tolerance breadth, and fecundity) and abiotic parameters (magnitude of temperature heterogeneity over space and temperature stochasticity over time), we determine which types of species are likely to persist in which types of environments. We simulate the management actions of increasing heterogeneity or decreasing stochasticity to quantify their effect on persistence likelihood and population size in a changing climate.

## Methods

To understand how spatio-temporal variation affects species’ abilities to track climate change along a temperature gradient, we built a stochastic metapopulation model simulating a species dispersing across an environment with changing temperatures (adapted from Backus & Baskett 2021; Backus et al. 2022). We represent a species as a discrete population over a series of connected patches on a one-dimensional temperature gradient (e.g., latitudinal or elevational gradient). To represent spatio-temporal heterogeneity in temperature, each patch contains several subpatches with unique temperatures that change stochastically over time at a designated level of variability. During each time step, the population cycles through reproduction, dispersal, and density-dependent survival. Species differ by dispersal (*γ*), thermal tolerance breadth (*σ*), and fecundity (*ρ*). Each environment is defined by the standard deviation in interannual environmental stochasticity (*S*) and the standard deviation in local climate heterogeneity (*H*). We randomized these five parameters for each simulation to explore how each influences species’ persistence and population size. We then implemented changing spatial heterogeneity or environmental stochasticity to simulate management actions that restore refugia.

### Environmental structure

We represent local temperature variation across space with the local climate heterogeneity parameter, *H* (Fig. 1e). Space in this model is a one-dimensional temperature gradient of *L* patches, representing large-scale latitudinal or elevational change (Urban et al. 2012). Each patch *x* contains *W* subpatches, representing small-scale variability in microclimates without an explicit spatial structure. Local subpatch (*w*) temperature is *T* (*x, w, t*) with a mean patch temperature of 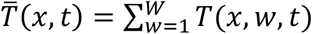 at time *t*. We set the local climate heterogeneity so that each patch has a standard deviation in local temperatures of

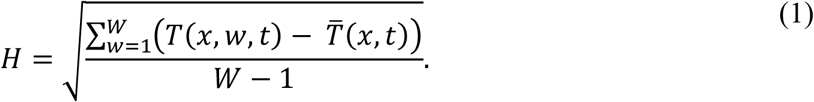

We represent temperature variation over time with the environmental stochasticity parameter, *S* (Fig. 1d). Each time step, all patches change in temperature by an average value of *τ*, with a stochastic component with autocorrelation κ, and standard deviation *S* around white noise *ω*(*t*). The stochastic component of yearly temperature change is 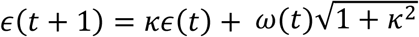, with the square root term to remove the effect of autocorrelation on the variance (Wichmann et al. 2005). Altogether, the temperature in patch *x*, subpatch *w*, changes over time is

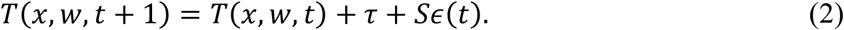

### Population dynamics

Simulated species have a population size of *n*(*x, w, t*) individuals in patch *x*, subpatch *w*, at time *t*. All individuals reproduce simultaneously at the beginning of each time step with a reproductive output *b*(*T*(*x, w, t*)) as a function of time- and location-dependent temperature *T*(*x, w, t*) (Fig. 1a). The ecological performance of many species, especially ectotherms, is skew-normal depending on temperature, with peak performance at a thermal optimum, a gradual decrease below the optimum, and a sharp decrease above the optimum (Huey & Kingsolver 1989; Norberg 2004). Therefore, the species’ temperature-dependence is skew-normal, given skewness constant *λ* with the highest values around the thermal optimum *ζ* and a sharp decrease above *ζ*. Given thermal tolerance breadth *σ* and fecundity *ρ*, the reproductive output is

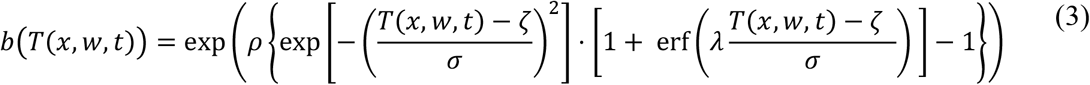

f(Urban et al. 2012). To incorporate demographic stochasticity, the number of propagules produced in patch *x*, subpatch *w* is a Poisson random variable with mean equal to *b*(*T*(*x, w, t*)), or *n*\*(*x, w, t*)∼Poisson (*n*(*x, w, t*) *b*(*T*(*x, w, t*))) (Melbourne & Hastings 2008).

Between reproduction and density-dependence, each propagule disperses from its origin (Fig. 1b), representing typical life histories of plants and many marine invertebrates. The model pools all propagules in a patch prior to dispersal, such that the total number of propagules in patch *x* at time *t* is 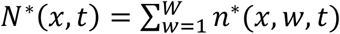. We adapt the heavy-tailed Laplace dispersal kernel to a discrete space analog, as most studies that fit empirical dispersal data to theoretical dispersal kernels show that heavy-tailed kernels (with accelerating spreading rates) outperform thinner-tailed kernels among a wide variety of species (Nathan et al. 2012). We define *γ* as the mean absolute distance (in patches) that individuals move and define the kernel parameter 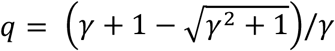. Thus, the probability of moving from patch *x* to patch *y* is

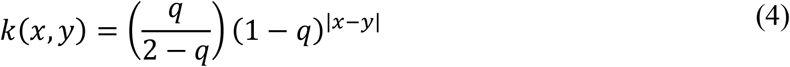

(Backus & Baskett 2021; Backus et al. 2022). All propagules disperse from patch *x* throughout all patches with a multinomial random vector. After arriving at patch *y*, propagules randomly distribute among the *W* subpatches of patch *y*. The resulting number of dispersed propagules in patch *y*, subpatch *w*, at time *t* is *n*\*\*(*y, w, t*).

Lastly, dispersed propagules compete for limited space and resources within each subpatch, given constant carrying capacity *K* in each subpatch (Fig. 1c). For simplicity, density-dependent survival is a variation on lottery competition (Sale 1978), where each individual has an equal probability of surviving, based on the Beverton-Holt density-dependence,

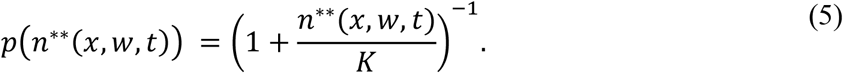

The total number of individuals that survive in patch *x*, subpatch *w*, after competition is a binomial random variable *n*(*x, t* + 1)∼Biomial (*n*\*\*(*x, w, t*), *p*(*n*\*\*(*x, w, t*))) (Melbourne & Hastings 2008). Though temperature does not affect density-dependent survival, this is immediately followed by reproduction that incorporates temperature-dependence (Eq. 3).

### Parameterization and implementation

For all simulations, the number total number of patches was *L* = 512 and the number of subpatches per patch was *W* = 8 (giving a total of 2^12^ discrete locations). The initial mean temperature across the temperature gradient varied linearly from 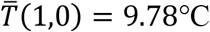 on the poleward edge to 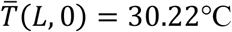 on the equatorward edge, though the species would only occupy a subset of this range. Annual temporal autocorrelation was *κ* = 0.767 based on the measured combined global land-surface air and sea-surface water temperature anomalies from 1880 to 1979 (GISTEMP Team 2019; Lenssen et al. 2019). The carrying capacity was a temperature-independent constant *K* = 8 per subpatch so that each patch could carry a total of 64 individuals at carrying capacity. In all simulations, we set the species’ thermal optimum to *ζ* = 20°C with a skewness *λ* = −2.7 to ensure that species performance is greatest at the center of the initial temperature gradient.

We ran two sets of simulations with 10^9^ replicates to quantify species persistence through gradually increasing temperatures under different species demographic values and degrees of spatio-temporal environmental variation. For each simulation, we generated a randomized environment and temporal time series based on the local temperature heterogeneity (*H*) and environmental stochasticity (*S*), each drawn from uniform distributions (Table 1). We generated species by drawing the mean dispersal distance (*γ*) from a log-uniform distribution and the thermal tolerance breadth (*σ*) and fecundity (*ρ*) from uniform distributions (Table 1).

**Table 1:** Parameter definitions and values used throughout simulations.

| Parameter | Symbol | Values | Units |
| --- | --- | --- | --- |
| Species' mean dispersal distance | $\gamma$ | Log-uniform; $10^{-4}$ to 1 | patches |
| Species' thermal tolerance breadth | $\sigma$ | Uniform; 2 to 5 | $^{\circ}\text{C}$ |
| Species' fecundity | $\rho$ | Uniform; 1.5 to 6 | - |
| Local temperature heterogeneity | $H$ | Uniform; 0 to 2 | $^{\circ}\text{C}$ |
| Environmental stochasticity | $S$ | Uniform; 0 to 1 | $^{\circ}\text{C}$ |
| Species' thermal optimum | $\zeta$ | 20 | $^{\circ}\text{C}$ |
| Skewness constant | $\lambda$ | -2.7 | - |
| Total patches | $L$ | 512 | patches |
| Subpatches per patch | $W$ | 8 | subpatches |
| Subpatch carrying capacity | $K$ | 8 | individuals |
| Mean annual temperature change | $\tau$ | 0 or 0.04 | $^{\circ}\text{C}/\text{year}$ |
| Annual temporal autocorrelation | $\kappa$ | 0.767 | - |

In the first set of simulations, we evaluated which types of species and environments were associated with species persistence during climate change without management action. To initialize the population size and range given the species’ biotic parameter values (*γ, σ*, and *ρ*) and the environmental parameter values (*H* and *S*), we placed 4 individuals on each subpatch and simulated the model for 200 years with no average temperature change, *τ* = 0°C per year. Then we simulated climate change by adjusting the temperature change to *τ* = 0.02°C per year reflecting intermediate emissions climate scenario (2°C over 100 years) and *τ* = 0.04°C per year reflecting a high emissions scenario (4°C over 100 years) (Urban et al. 2012, IPCC 2014). We tracked whether or not the species persisted anywhere the landscape, and, following the methodology of a global sensitivity analysis (Cariboni et al. 2007), we ran a random forest classification (randomForest 4.6-14 package, R Version 4.03) with persistence/extinction as the dependent variable and *γ, σ, r, S*, and *H* as the dependent variables.

In the second set of simulations, we evaluated which types of species would benefit from conservation management actions of increasing the local heterogeneity or decreasing the environmental stochasticity. We used the same set parameters generated above, but we modified the initial local heterogeneity to a set value *H*_1_ = 1 and initialized the population again over 200 years with no average temperature change. Then we simulated how two management scenarios affected the final population after 100 years of increasing temperatures under both the 2°C and 4°C scenarios. When unmanaged, we left the local heterogeneity at *H*_2,*a*_ = *H*_1_ = 1, and when restoring heterogeneity, we changed the local heterogeneity to *H*_2,*b*_ = 2. We performed a similar set of simulations keeping heterogeneity constant but reducing stochasticity, such that the initial stochasticity was *S*_1_ = 0.5, and the two scenarios were *S*_2,*a*_ = 0.5 (unmanaged) and *S*_2,*b*_ = 0.25 (managed). In these simulations, we considered species to have benefited from restoration if the final population under management was greater than 105% of the value for the simulations without management. We compared population size rather than persistence to help us better detect potential negative effects of management and gain a more nuanced understanding of potential benefits.

## Results

Out of all species we simulated, 61.2% persisted without management intervention under 4°C of temperature increase over 100 years, and 91.3% persisted under 2°C of temperature increase. Under both climate change scenarios, simulated species were less likely to persist or had a lower population size if they had shorter mean dispersal distance *γ*, narrower thermal tolerance *σ*, or lower fecundity *ρ*, and when they were in environments with higher interannual variation in stochasticity *S* or lower local heterogeneity *H* (Fig. 2, S1, S2; random forest out-of-bag error: 6.87% under 4°C, 6.47% under 2°C). In general, species with multiple biological limitations (e.g., species with both short dispersal and narrow thermal tolerance) were less likely to persist through climate change (Fig. 3, S3).

**Figure 2:**
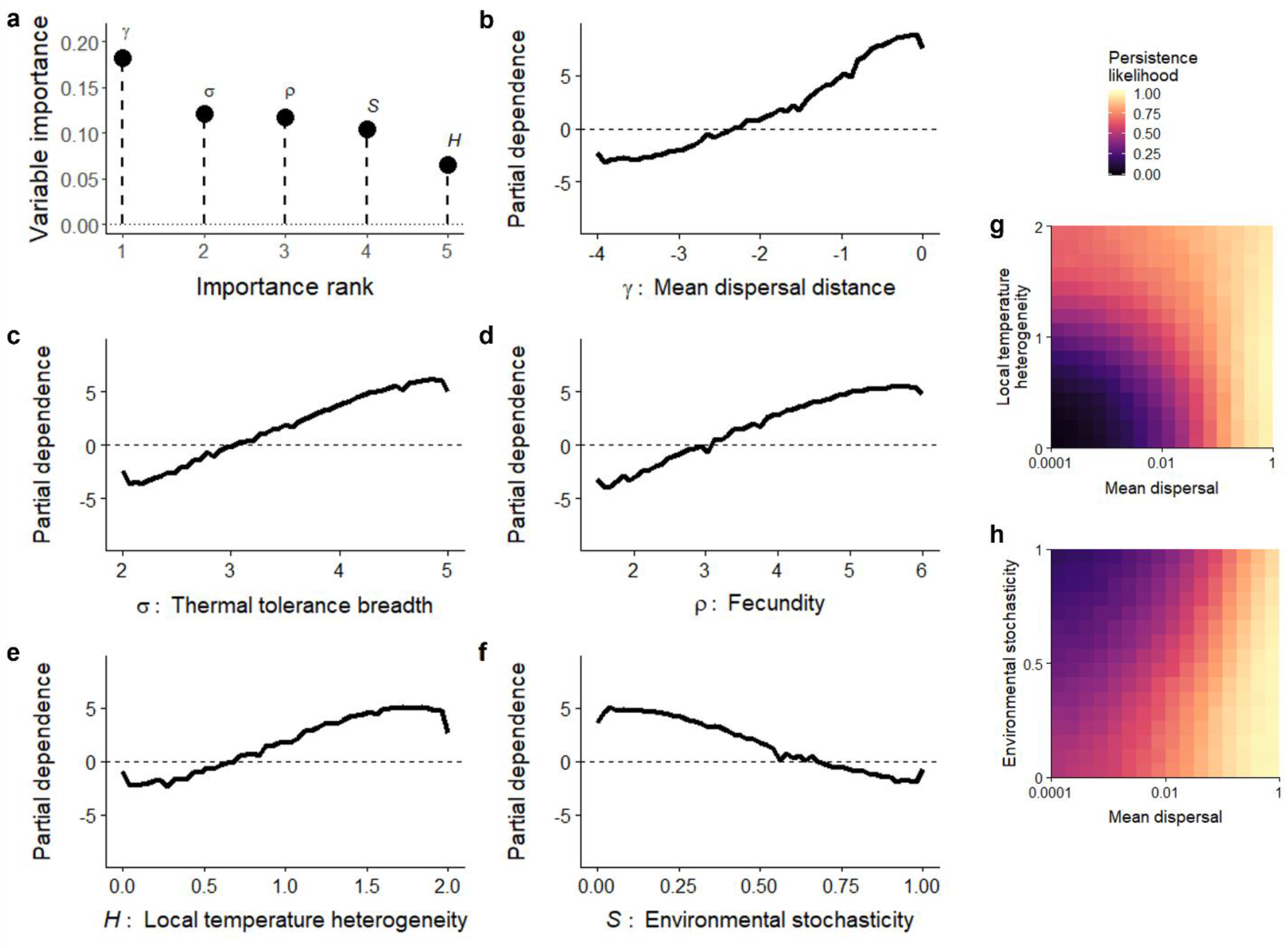
Importance of biotic and abiotic parameters from random forest classifications for randomized species persistence through climate change by 4°C over 100 years. (a) Relative unscaled permutation importance of dispersal distance (*γ*), thermal tolerance breadth (*σ*), fecundity (*ρ*), local climate heterogeneity (*H*), and environmental stochasticity (*S*). (b-f) Partial dependence of biotic and abiotic parameters. Vertical axis is the log-odds of species’ persistence through the simulation (higher values indicate a greater persistence likelihood). (g-h) Persistence likelihood grouped by mean dispersal ability *γ* (horizontal axis) and either local temperature heterogeneity *H* (vertical axis in g) or environmental stochasticity *S* (vertical axis in h).

**Figure 3:**
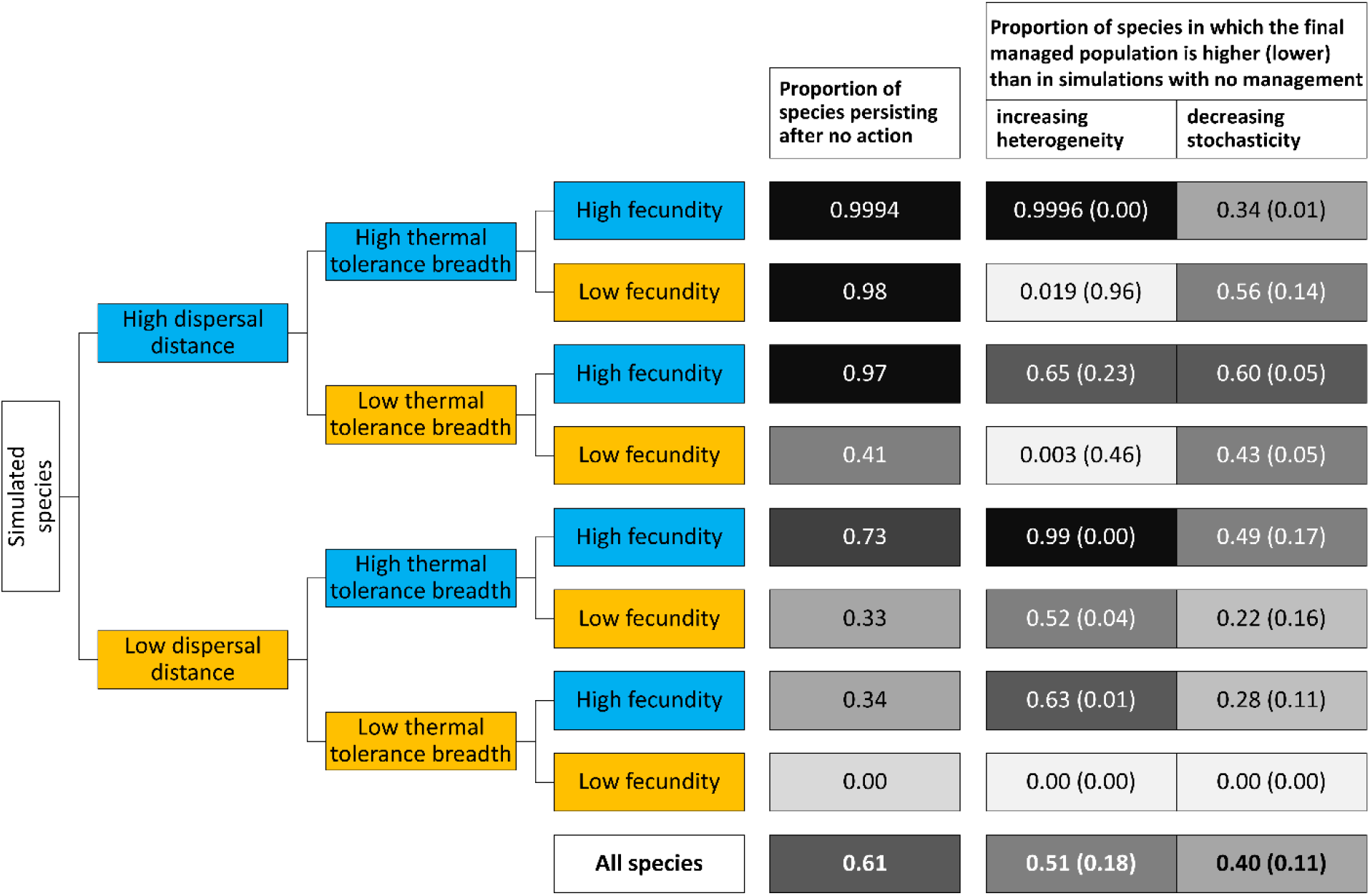
Results from simulations when the average temperature increases by 4°C over 100 years, categorizing species into highest or lowest quartiles of biotic parameters. Higher trait values (in the upper quartile) are highlighted in blue, and lower trait values are highlighted in orange. The final two columns represent the proportion of species that benefit from either increasing heterogeneity or decreasing stochasticity (final population size of the species more than 105% relative to simulations without management action) and the proportion that are negatively affected in parentheses (final population size less than 95% relative to simulations without management action). Darker colors in the final three columns represent higher values.

On average, restoring heterogeneity along the temperature gradient (by doubling *H*) improved a species’ likelihood of persisting through 4°C of temperature increase over 100 years more often than it decreased a species’ likelihood of persisting (Fig. 3, Fig. 4a,c,e). Restoring heterogeneity also generally increased a species’ population size relative to no action. However, when the temperature increased by only 2°C over 100 years, restoring local heterogeneity was more likely to decrease a species’ population size, and the species that were more likely to benefit were those that were already likely to persist (Fig. S4, S5). In simulations with 4°C of temperature increase, species with shorter average dispersal particularly benefited from increased heterogeneity (Fig. 3). Though many of the species with the shortest dispersal ranges went extinct regardless of management, increasing the local heterogeneity was more likely to prevent extinction and increase the final population size and rarely decreased population size or caused extinction (Fig. 4a). Species with longer dispersal ranges were less likely to go extinct without management, and increasing heterogeneity was less likely to benefit and more likely to decrease the population size or cause the extinction of these species. Species with narrow thermal tolerance or low fecundity were also unlikely to persist without heterogeneity restoration, but these species did not benefit as strongly and were more likely to experience negative effects from increased heterogeneity when compared with species with shorter dispersal ranges (Fig. 4c,e).

**Figure 4:**
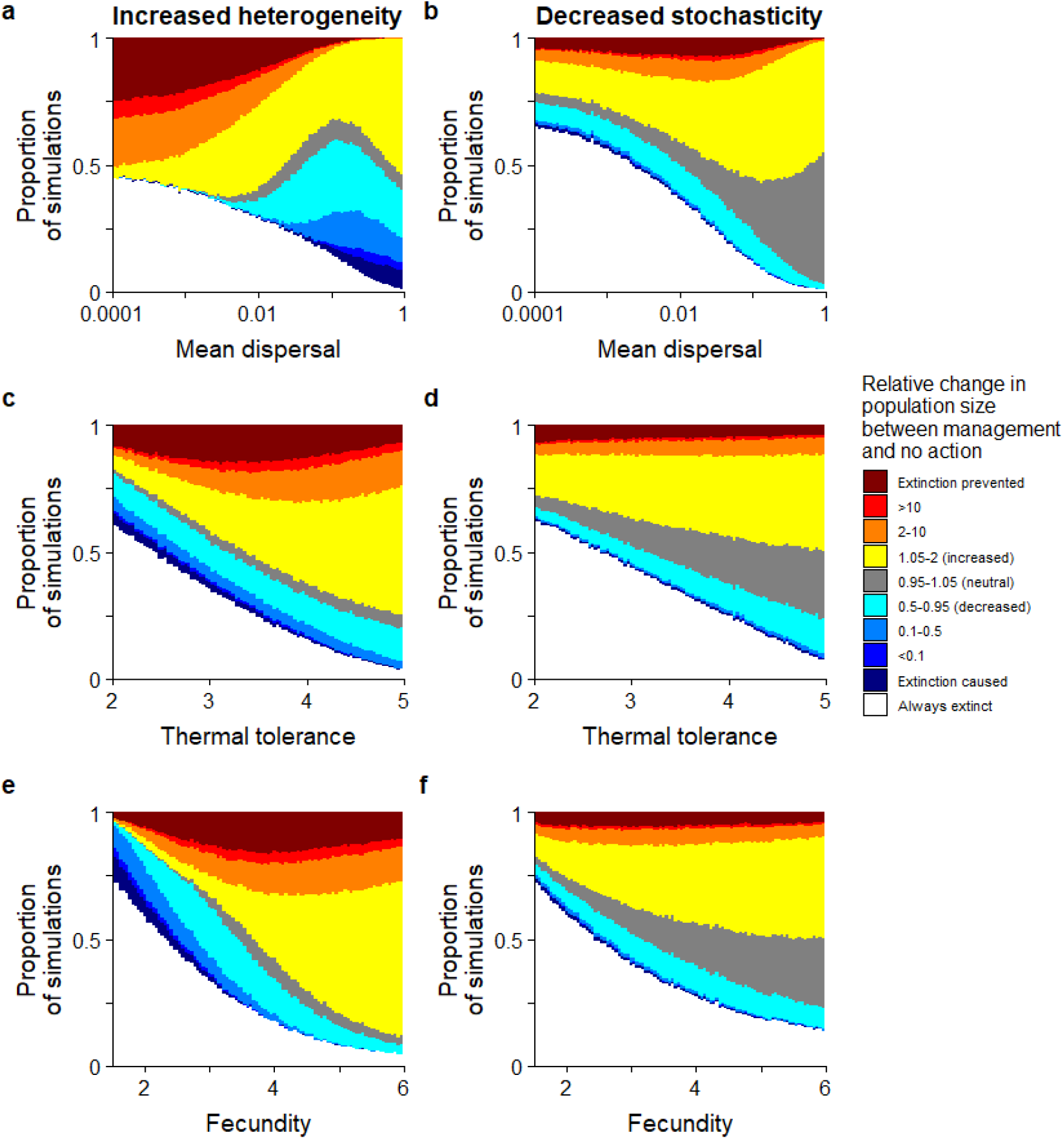
Relative effect of increasing local heterogeneity (a,c,e) or decreasing environmental stochasticity (b,d,f) on a species’ fate under a 4°C increase over 100 years compared to no action, under a range of values for mean dispersal *γ* (a,b), thermal tolerance (c,d), and fecundity (e,f). The vertical axis indicates frequencies of outcomes within bins that represent different species fates: white indicates simulations where the species goes extinct during climate change regardless of management. Dark red indicates simulations where a species goes extinct with no action but persists with management. Hues from yellow to red indicate simulations where management increases a species’ population size relative to no action. Gray indicates simulations where a species persists, and management has little effect on the final population size. Hues from cyan to blue indicate simulations where management decreases a species’ population size relative to no action. Dark blue indicates that management caused the species to go extinct.

Decreasing the environmental stochasticity along the temperature gradient (by halving the value of *S*) typically increased the persistence likelihood and population size (relative to no action) of species throughout our simulations under both the 2°C and 4°C increase scenarios (Fig. 4b,d,f, S5). Decreasing the stochasticity did not strongly benefit species with very short dispersal, narrow thermal tolerance, or low fecundity, as many of these species were likely to go extinct regardless of management action. Species with longer dispersal, narrower thermal tolerance, and higher fecundity were more likely to benefit from decreased stochasticity.

Comparing both restoration strategies in the 4°C increase scenario, we found that restoring heterogeneity was particularly beneficial to species with low dispersal (below the 25% quantile) but with high values in one or both of the other two traits (above the 75% quantile) (Fig. 3). For these combinations, restoring heterogeneity either prevented the species from going extinct or increased the final managed population size (at least 105% the population size of the unmanaged population) in a high proportion of simulations. However, species with high dispersal and low fecundity were unlikely to benefit from increased heterogeneity but were somewhat likely to benefit from decreased stochasticity. Species with low values in all three traits went extinct in every simulation, regardless of management.

## Discussion

Overall, we found that simulated species were more likely to persist through climate change when they lived in temperature gradients with higher habitat heterogeneity and lower environmental stochasticity. This arises because species in heterogeneous environments do not need to disperse as far to reach analogous during climate change, and species are more likely to persist with more consistent year-to-year changes of lower environmental stochasticity. We also find that many species with strong biological limitations (e.g. short distance, narrow thermal tolerance, or low fecundity) might benefit from conservation management strategies that increase local climate heterogeneity or reduce interannual variation in climatic conditions. Although some of the species we modeled were negatively affected by these restoration efforts, the majority of species experienced long-term benefits from restoring heterogeneity or decreasing stochasticity, especially in our more extreme climate change simulations. These results build on previous findings that climate and habitat heterogeneity can maintain biodiversity in the absence of climate change (Stein et al. 2014, Gámez-Virués et al. 2015), suggesting that heterogeneity can also create refugia-like conditions to buffer against climate change.

### Approaches to restoring refugia

Restoring climate refugia by restoring local climate heterogeneity might be achieved by modifying physical landscapes and ecological communities to resemble a pre-agricultural or pre-industrial state (Palmer et al. 2010; Morelli et al. 2016). In human-modified forest landscapes that have been converted to agriculture, this means not only maintaining several high-quality patches with diverse plant communities, but also scattering trees between the dense areas (Fischer et al. 2010; Arroyo-Rodríguez et al. 2020). Approaches to integrate natural habitat into human-dominated landscapes, such as wildlife-friendly framing practices (Green et al. 2005) and their incentivization through agri-environmental schemes (Donald & Evans 2006), provide an opportunity to restore lost heterogeneity (Fischer et al. 2008) that might then improve species persistence through climate change. For example, many dispersal-limited climate-threatened herpetofauna in Romania could lose all livable climate space by the 2050s (Popescu et al. 2013). However, because of recent political change in eastern Europe, many areas that were previously deforested for agriculture have become abandoned and fragmented (Bălteanu & Popovici 2010). Others have already suggested restoring these abandoned areas into traditional farming landscapes with heterogeneous vegetation to benefit the people and biodiversity in the region (Fischer et al. 2012), and our theoretical results suggest this could also help buffer dispersal-limited species persist through rapid climate change.

Restoring climate refugia by reducing environmental stochasticity might be difficult to manage on a large scale, but it might be possible in a few situations. Canopy cover can affect temperature extremes, acting as climate refugia on forest floors and riparian environments (Davies 2010; De Frenne et al. 2013; Reiter et al. 2020). Clear-cut areas experience a greater range in temperatures than deciduous or mixed forests (Barbier et al. 2008). Forestry practices that promote diverse stands of deciduous or mixed trees could minimize the magnitude of temperature extremes and minimize stochasticity, which our theoretical results suggest could increase the persistence of long-distance-dispersing, low-fecundity species under climate change.

### Species traits associated with benefits from refugia restoration

Species with limited dispersal ability are expected to be among the most at risk of extinction from climate change because they are unlikely to disperse far enough to reach analogous climates to their historic ranges given projected rates of climate change (Pearson 2006; Urban 2015). Accordingly, we found that the shortest-distance dispersing species were most likely to go extinct when unmanaged during our climate change simulations, and these species disproportionately benefited from simulated heterogeneity restoration. However, heterogeneity restoration only benefited dispersal-limited species if they also had either broad thermal tolerance or high fecundity, while species that were limited by all three demographic traits went extinct regardless of management intervention. Given these expectations, example candidates of species likely to benefit from restoring habitat heterogeneity might include plant species that would be classified as ruderal or stress tolerant (Grime 1977), such as early-to-mid successional plants (Meier et al. 2012) or alpine shrubs adapted to unique soil types (Damschen et al. 2012). Similarly, many reptiles and amphibians have limited dispersal ability and narrow climate tolerance, but relatively high fecundity (Araújo et al. 2006), suggesting that they might especially benefit from heterogeneity restoration.

As compared to restoration of spatial heterogeneity restoration, reducing the environmental stochasticity was more likely to benefit species with longer average dispersal and lower fecundity in our simulations. However, many of the species that disproportionally benefitted from stochasticity reduction were likely to persist through climate change even without restoration. Moreover, long-distance-dispersing, low-fecundity species might be rare given the typical positive correlation between dispersal distance and fecundity in taxon such as plants (Beckman et al. 2017). Because dispersal allows individuals to reach higher quality environments under temporally unpredictable conditions, it can drive the evolution of dispersal under high environmental stochasticity (Levin et al 1984; McPeek & Holt 1992), and species with greater dispersal ability could be less common in historically low-stochasticity environments. Species with high dispersal and low fecundity could be large longer-lived animal species with high investment in offspring care, including many mammals and birds, though longer life histories also provide a buffer against environmental stochasticity (Halley et al. 2018). Even birds with higher dispersal ability have become uncoupled with their optimal climate in the past few decades (Viana & Chase 2022), but this could be driven by transient populations in heterogenous environments (Coyle et al. 2013).

Restoration to increase spatial heterogeneity or buffer environmental stochasticity did not alleviate extinction risk for species within the bottom quartiles for all three of dispersal, climate tolerance, and fecundity in our simulations. For such species, alternative management approaches to mitigating climate-driven extinction (Heller & Zavaleta 2009; Backus et al. 2022) might more effectively reduce extinction risk. For example, assisted migration, the intentional movement of the species to a location that is projected to be more suitable under projected climate change (McLachlan et al. 2007; Schwartz et al. 2012), could allow a species to reach a suitable future climate despite dispersal and fecundity limitations, although moving a species into a new location could bring the risks of invasion (Mueller & Hellmann 2008; Hewitt et al. 2011). Another risk of assisted migration is that these species might fail to establish in their recipient communities after relocation because of species interactions, especially for species with low thermal tolerance in a stochastic environment, causing a loss to the source population (Peterson & Bode 2021; Backus & Baskett 2021). For such species, one potential management strategy to promote thermal tolerance is assisted gene flow, the introduction of climate-tolerant traits or other beneficial traits from other populations into climate-threatened populations, such as through the relocation of locally adapted individuals within a species’ range (Aitken & Whitlock 2013). Furthermore, for species with sufficient dispersal to track climate change but are limited by habitat fragmentation, management to increase connectivity between high-quality patches, such as the establishment of corridors, might prove more effective (Heller & Zavaleta 2009; Lawler & Olden 2011).

### Model assumptions and future directions

Our model provides a first step towards quantifying the expected efficacy of restoring climate refugia, a widely considered conservation tool among several types of ecosystems (Davies 2010; Chollett & Mumby 2013; Morelli et al. 2016). Further theoretical research might expand on many of the simplifying assumptions we made in this model. In particular, our model considers single populations in isolation of an ecological community, but a species’ ability to shift its range with climate change generally depends on species interactions (Urban et al. 2012). These interactions could be crucial, as competing species could slow climate tracking by preventing species from dispersing poleward, especially if there is differential dispersal ability between interacting species (Urban et al. 2012). If longer-distance dispersing species colonize newly restored environments before shorter-distance dispersing species, competition could dampen the benefits of heterogeneity restoration. Also, our model assumed a constant carrying capacity, reflecting equal habitat quality throughout our simulated landscape. Patchy environments could mean that short-dispersing species would be unable to disperse over larger scale temperature gradients (e.g., latitudinally), which could increase the benefit of restoring local climate heterogeneity. As managers continue to consider climate refugia as a conservation tool for climate change, further ecological models adapted to specific species and ecosystems can reveal potential costs and benefits of management actions that affect climate heterogeneity and environmental stochasticity in practice.

## Acknowledgements

We would like to thank L. Bay, R. Gates, S. Harrison, C. Logan, M. McClure, C. Muhlfeld, S. Sawyer, M. Schwartz, R. Waples, and A. Whipple for their thoughtful conversations at the managed relocation workshop at UC Davis that informed this manuscript. This project was funded by the National Science Foundation, grant #1655475.

