## Supplemental figures for "Restoring local climate refugia to enhance the capacity for dispersal-limited species to track climate change"

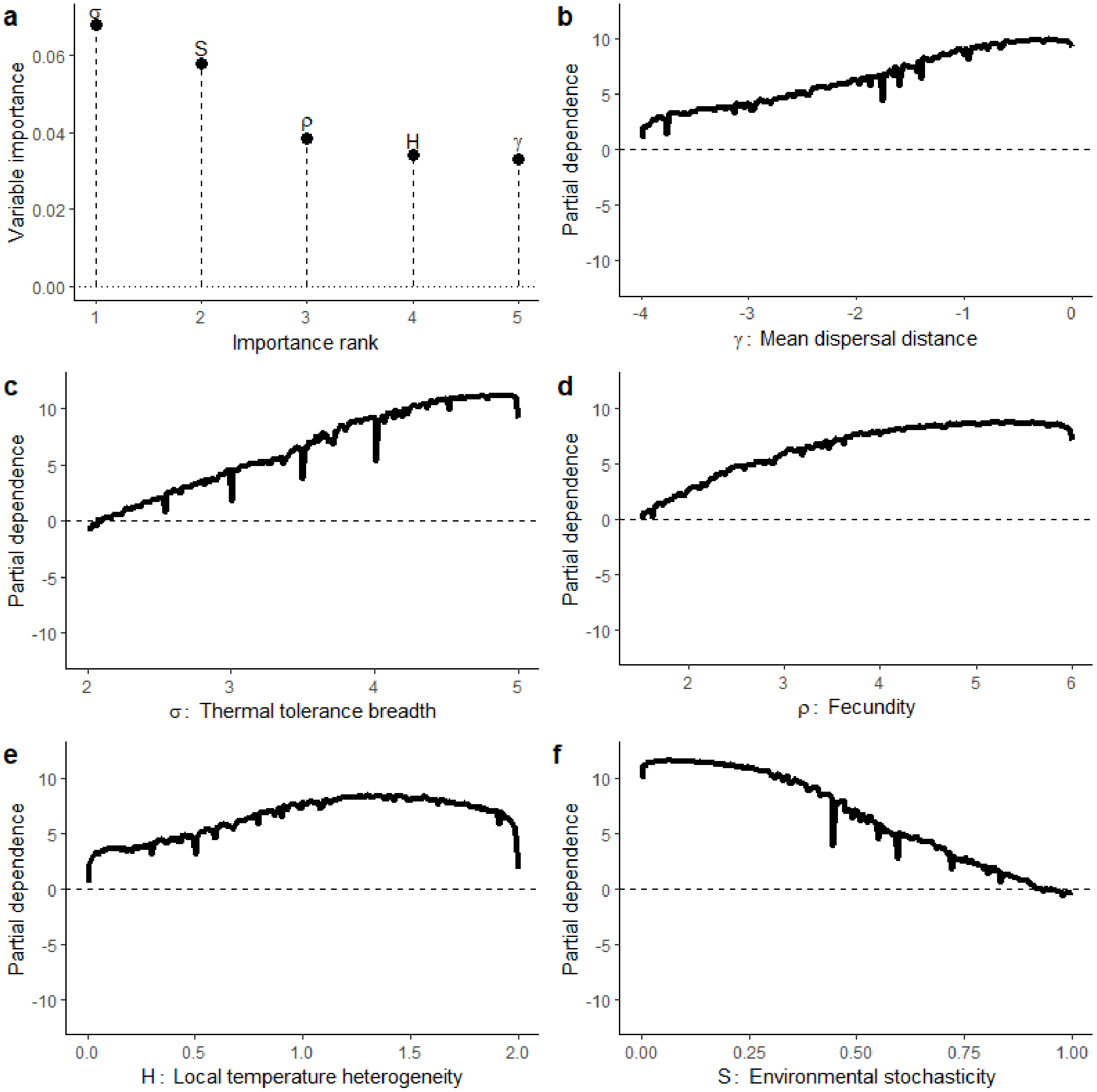


**Figure S1**: Importance of biotic and abiotic parameters from random forest classifications for randomized species persistence through climate change simulations with an increase of 2°C over 100 years. (a) Relative unscaled permutation importance of dispersal distance ($\gamma$), thermal tolerance breadth ($\sigma$), fecundity ($\rho$), local climate heterogeneity ($H$), and environmental stochasticity ($S$). (b-f) Partial dependence of the values of the species and environmental trait values. The y-axis is the log-odds of whether the species persists through the simulation (higher values indicate a greater persistence likelihood).

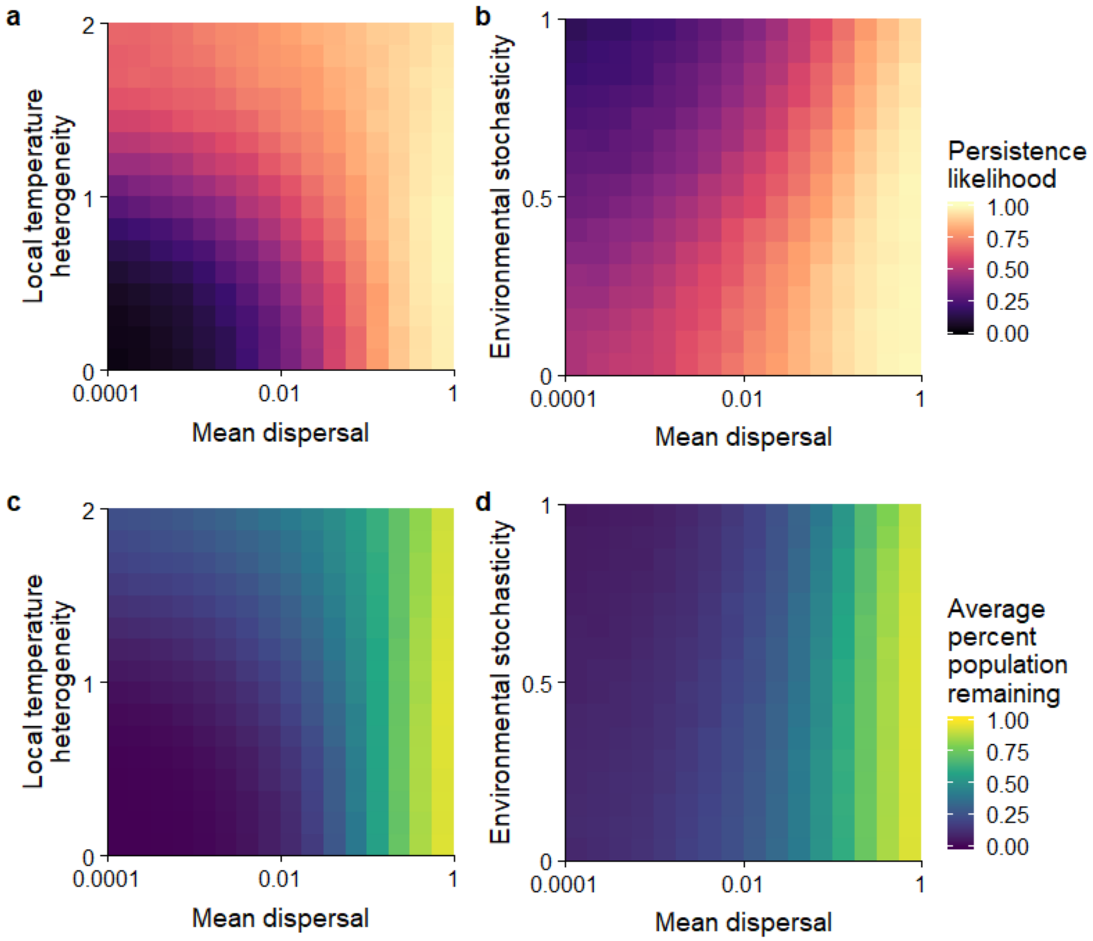


**Figure S2**: Average simulation outcomes for a species experiencing an increase of 4°C over 100 years of climate change without restoration. (a, b) Persistence likelihood averaged across species with randomly drawn values for demographic traits and spatio-temporal environmental variability, grouped by mean dispersal ability $\gamma$ (horizontal axis) and either local temperature heterogeneity H (vertical axis in a) or environmental stochasticity S (vertical axis in b). (c-d) Average percent of the initial population size remaining after climate change for randomized species grouped by mean dispersal ability $\gamma$ (horizontal axis) and local temperature heterogeneity H (vertical axis in c) or environmental stochasticity S (vertical axis in d).

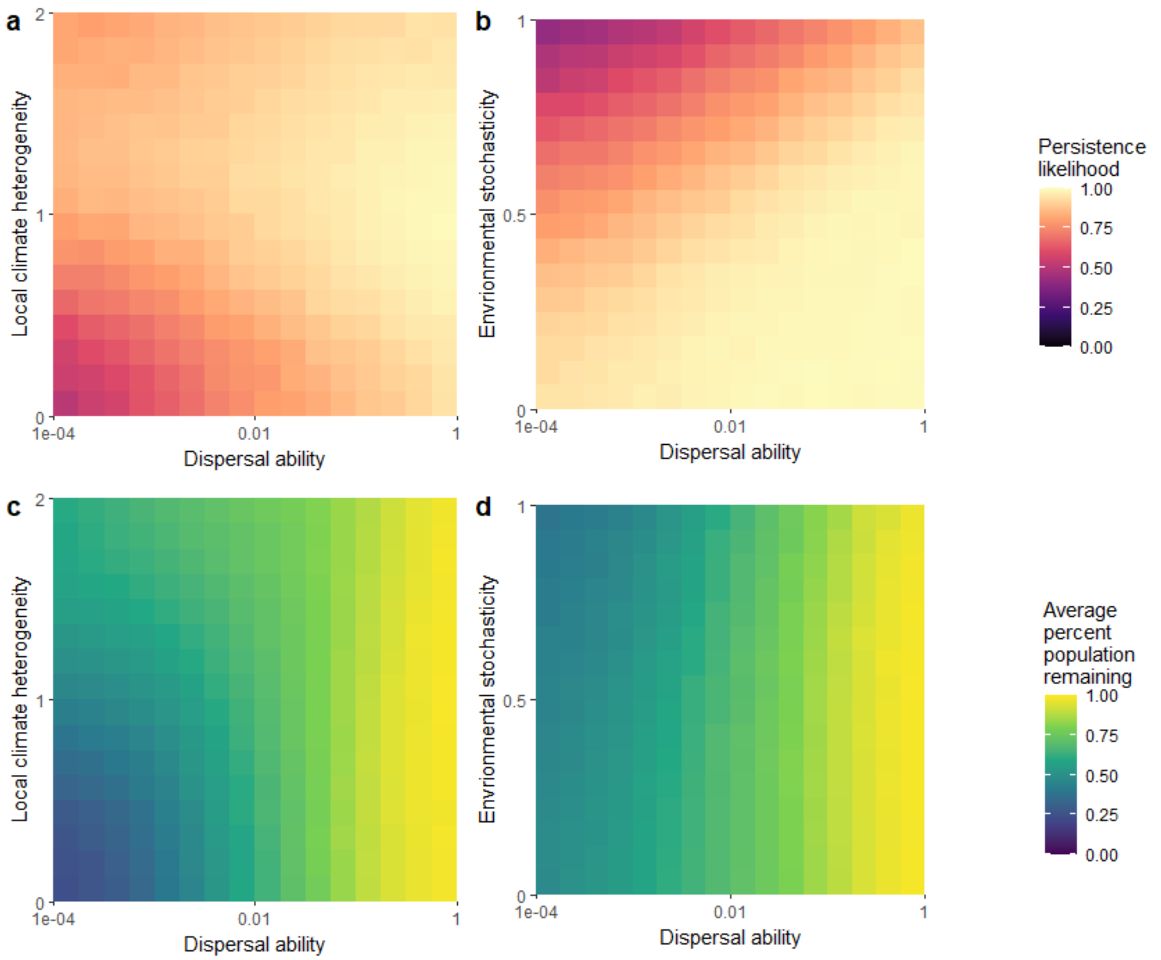


**Figure S3**: Average simulation outcomes for a species experiencing an increase of 2°C over 100 years of climate change without restoration. (a, b) Persistence likelihood averaged across species with randomly drawn values for demographic traits and spatio-temporal environmental variability, grouped by mean dispersal ability γ (horizontal axis) and either local temperature heterogeneity H (vertical axis in a) or environmental stochasticity S (vertical axis in b). (c-d) Average percent of the initial population size remaining after climate change for randomized species grouped by mean dispersal ability γ (horizontal axis) and local temperature heterogeneity H (vertical axis in c) or environmental stochasticity S (vertical axis in d).

| Simulated species | | | Proportion of species persisting after no action | Proportion of species in which the final managed population is higher (lower) than in simulations with no management: increasing heterogeneity | Proportion of species in which the final managed population is higher (lower) than in simulations with no management: decreasing stochasticity |
|---|---|---|---|---|---|
| High dispersal distance | High thermal tolerance breadth | High fecundity | 0.9998 | 1.00 (0.00) | 0.35 (0.0008) |
| | | Low fecundity | 0.9997 | 0.007 (0.97) | 0.51 (0.03) |
| | Low thermal tolerance breadth | High fecundity | 0.99 | 0.85 (0.06) | 0.62 (0.02) |
| | | Low fecundity | 0.84 | 0.001 (0.92) | 0.76 (0.06) |
| Low dispersal distance | High thermal tolerance breadth | High fecundity | 0.98 | 0.98 (0.004) | 0.59 (0.06) |
| | | Low fecundity | 0.95 | 0.09 (0.87) | 0.63 (0.18) |
| | Low thermal tolerance breadth | High fecundity | 0.81 | 0.33 (0.54) | 0.70 (0.07) |
| | | Low fecundity | 0.23 | 0.00 (0.53) | 0.13 (0.04) |
| | | **All species** | **0.91** | **0.38 (0.48)** | **0.61 (0.07)** |

**Figure S4**: Results from simulations when the average temperature increases by 2°C over 100 years, categorizing species into highest or lowest quartiles of biotic parameters. Higher trait values (in the upper quartile) are highlighted in blue, and lower trait values are highlighted in orange. The final two columns represent the proportion of species that benefit from either increasing heterogeneity or decreasing stochasticity (final population size of the species more than 105% relative to simulations without management action) and the proportion that are negatively affected in parentheses (final population size less than 95% relative to simulations without management action). Darker colors in the final three columns represent higher values

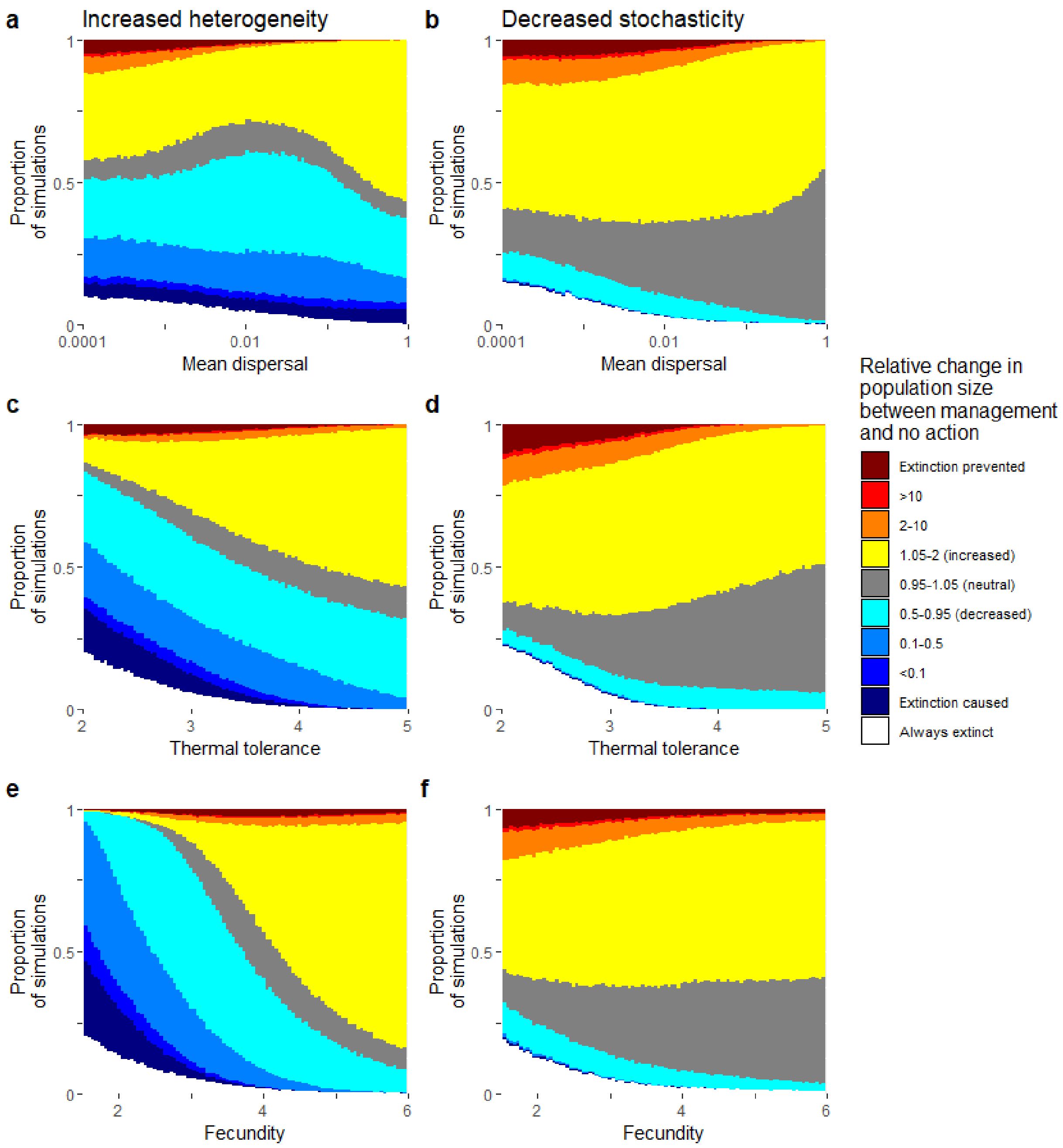


**Figure S5**: Relative effect of increasing local heterogeneity (a,c,e) or decreasing environmental stochasticity (b,d,f) on a species' fate during an increase of 2°C over 100 years of climate change compared to no action, under a range of values for mean dispersal $\gamma$ (a,b), thermal tolerance (c,d), and fecundity (e,f). The vertical axis indicates frequencies of outcomes within bins that represent different species fates: white indicates simulations where the species goes extinct during climate change regardless of whether there is management action. Dark red indicates simulations where a species goes extinct during climate change with no action but persists with management action. Hues from yellow to red indicate simulations where management increases a species' population size relative to no action. Gray indicates simulations where a species persists regardless of management action, but management has little effect on the final population size. Hues from cyan to blue indicate simulations where management decreases a species' population size relative to no action. Dark blue indicates simulations where management caused the species to go extinct where the species would have persisted with no action.
